# Dysregulation in mTOR/HIF-1 signaling identified by proteo-transcriptomics of SARS-CoV-2 infected cells

**DOI:** 10.1101/2020.04.30.070383

**Authors:** Sofia Appelberg, Soham Gupta, Anoop T Ambikan, Flora Mikaeloff, Ákos Végvári, Sara Svensson Akusjärvi, Rui Benfeitas, Maike Sperk, Marie Ståhlberg, Shuba Krishnan, Kamal Singh, Josef M. Penninger, Ali Mirazimi, Ujjwal Neogi

**Affiliations:** Public Health Agency of Sweden, Solna, Sweden; Division of Clinical Microbiology, Department of Laboratory Medicine, Karolinska Institute, ANA Futura, Campus Flemingsberg, Stockholm, Sweden; Division of Chemistry I, Department of Medical Biochemistry and Biophysics, Karolinska Institutet, Stockholm, Sweden; National Bioinformatics Infrastructure Sweden (NBIS), Science for Life Laboratory, Department of Biochemistry and Biophysics, Stockholm University, S-10691 Stockholm, Sweden; Department of Molecular Microbiology and Immunology and the Bond Life Science Center, University of Missouri, Columbia, MO 65211, USA; Institute of Molecular Biotechnology of the Austrian Academy of Sciences, Dr. Bohr-Gasse 3, 1030 Vienna, Austria; Department of Medical Genetics, Life Science Institute, University of British Columbia, Vancouver, V6T 1Z3, British Columbia, Canada; National Veterinary Institute, Uppsala, Sweden

## Abstract

How Severe Acute Respiratory Syndrome Coronavirus-2 (SARS-CoV-2) infections engage cellular host pathways and innate immunity in infected cells remain largely elusive. We performed an integrative proteo-transcriptomics analysis in SARS-CoV-2 infected HuH7 cells to map the cellular response to the invading virus over time. We identified four pathways, ErbB, HIF-1, mTOR and TNF signaling, among others that were markedly modulated during the course of the SARS-CoV-2 infection *in vitro*. Western blot validation of the downstream effector molecules of these pathways revealed a significant reduction in activated S6K1 and 4E-BP1 at 72 hours post infection. Unlike other human respiratory viruses, we found a significant inhibition of HIF-1α through the entire time course of the infection, suggesting a crosstalk between the SARS-CoV-2 and the mTOR/HIF-1 signaling. Further investigations are required to better understand the molecular sequelae in order to guide potential therapy in the management of severe COVID-19 patients.

## Introduction

The recent emergence of the coronavirus disease (COVID-19) pandemic caused by Severe Acute Respiratory Syndrome Coronavirus-2 (SARS-CoV-2) has created a public health emergency across the globe ^1–3^. SARS-CoV-2, a single-stranded positive-sense RNA virus, is the seventh coronavirus that infects humans and belongs to the β-coronavirus family. Due to limited knowledge on molecular mechanisms of infection and pathogenesis there is currently no available vaccine or specific therapeutics to treat or prevent SARS-CoV-2 infection.

Understanding the viral dynamics and host responses to the virus are necessary to design better therapeutic strategies for COVID-19 patients. Within the short period of the pandemic, there are few reports (pre-prints) on different levels of omics data (transcriptomics, proteomics, and metabolomics) from cell culture infected with SARS-CoV-2 as well as patient material that aimed to elucidate potential mechanisms of the host immune response and disease pathogenesis of SARS-CoV-2. However, the steady state measurements fail to reveal the dynamic changes of the host and viral proteins during the course of the infection. Thus, the temporal changes in gene expression and protein synthesis in different phase of the infection has not yet been reported.

To provide a comprehensive assessment of the cellular response to SARS-CoV-2, we performed a time series integrative proteo-transcriptomics analysis in infected HuH7 cells ranging from the early phase of infection until the virus reached the ability to initiate cytopathic effect at ~72 hours post infection (hpi).

## Results

### Dynamics of the SARS-CoV-2 infection in HuH7 cell lines

HuH7 cells were infected with SARS-CoV-2 at MOI 1.0 ^4^. Cells were collected at 24hpi, 48hpi and 72hpi and subjected to viral RNA quantification by quantitative PCR, transcriptomics by Illumina NextSeq550 and proteomics by tandem tag labeled mass spectrometry (TMT-MS). Transcriptomics, proteomics and proteo-transcriptomics data were further analyzed using in-depth bioinformatics (Fig 1a). qPCR targeting the E (envelope) gene of the SARS-CoV-2 identified a gradual increase in cellular viral RNA over time (p<0.05, repeated measure ANOVA) (Fig 1b). RNAseq analysis detected viral RNA at all the time points; 24hpi, 48hpi and 72hpi (Fig 1c). The TMT-based quantitative proteomics also identified statistically significant increase (p<0.05, repeated measure ANOVA) in SARS-CoV-2 proteins nucleocapsid (N), Membrane (M) and Spike (S) over time (Fig 1d). Thus, SARS-CoV-2 exposure of HuH7 cells results in effective infections that over time lead to enhanced viral RNA and viral protein production, required to assemble viral progeny.

**Figure 1.**
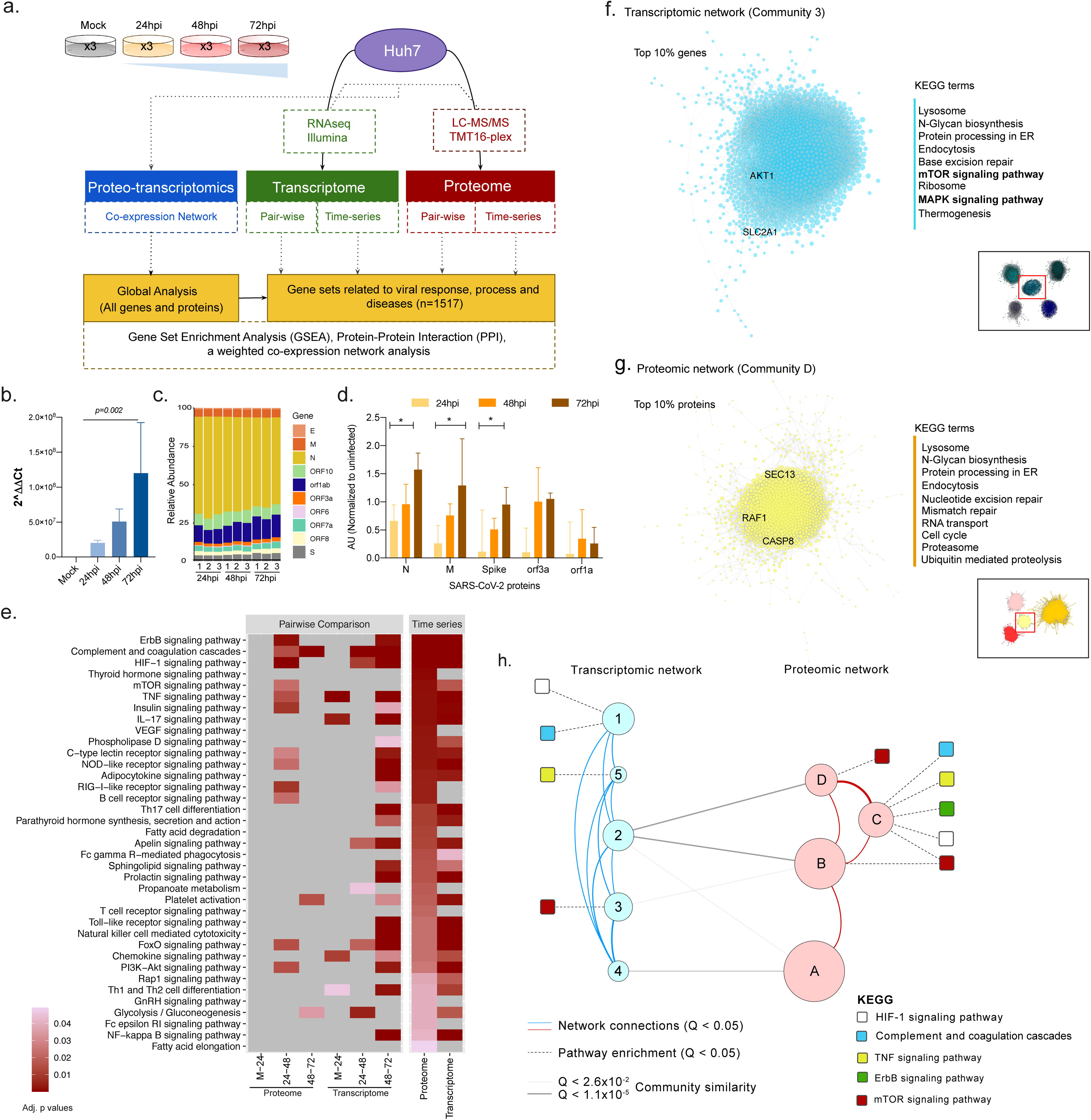
**(a)** Brief methodology. **(b)** Viral RNA quantification using qPCR targeting the E gene of SARS-CoV-2. **(c)** Detected viral genes and open reading frame in the RNAseq experiment. **(d)** Temporal dynamics of detected proteins in the HuH7 cells by TMT-MS. **(e)** Gene set enrichment analysis using the genes related to viral response, process and diseases in single omics level by pairwise comparative analysis and time series analysis at individual omics level. Significant (adjusted p values) KEGG terms enriched for upregulated genes were represented as heatmap. The lower adjusted p values are shown in dark red color and higher ones with light red color, non-significant pathways are represented in grey color. **(f-g)** Network analysis using genes and proteins. Analysis of the most central communities in each network highlights key KEGG terms (right) among the top 10% associated genes and proteins. The top 10% correlations (Spearman rho > 0.95, FDR < 0.05) were selected in the most central community in transcriptomic **(f)** and proteomic **(g)** networks (inset) based on mean normalized degree. The top KEGG terms associated with each of the two communities (FDR < 0.05) are highlighted, as well as genes that had been previously found in Fig 1e. **(h).** A proteo-transcriptomic network analysis highlights coordinated expression and functional changes in response to viral infection. Communities (circles) in transcriptomic and proteomic networks, where node size is proportion to the number of elements (728 - 2519). Edges indicate association (Q<0.05) with KEGG terms (dashed), network edges (solid red and blue), or community similarity (solid gray).

#### Host cellular response against SARS-CoV2 *in vitro*

After having established effective SARS-CoV-2 infections, we assessed the cellular host response to the virus infection. We found that 2622 genes and 1819 proteins were increased whereas 2856 genes and 1743 proteins were decreased significantly (false discovery rate <0.05) over the time despite distinct coverage (19997 protein-coding genes vs 7757 proteins quantified). We next performed gene set enrichment analyses using the differentially expressed genes/proteins that are related to viral response, process and diseases (targeted analysis) obtained from Gene Ontology (GO), REACTOME and “Rare_Diseases_AutoRIF_Gene_Lists” library and mapped to KEGG (Human_2019) terms. Figure 1e shows a heatmap of significantly enriched KEGG terms that are dysregulated in both our proteomics and transcriptomics analysis using for pairwise and time series analysis in uninfected and SARS-CoV-2 infected HuH7 cells. Of note, the downregulated genes did not identify any KEGG term with adjusted p value <0.05. Among the most significantly upregulated pathways, mining both proteomics and transcriptomics data, were pathways associated with cell proliferation and apoptosis, such as ErbB, PI3K-Akt, HIF-1, and mTOR signaling, and pathways that are related to innate immune responses such as TNF, NOD-like receptor (NLR) and RIG-I signaling (Fig. 1e). In addition, we observed upregulated platelet activation, complement cascades, FOXO signaling, and glycolysis (Fig. 1e), indicating that the SARS-CoV-2 infection induce pathways linked to thrombosis and metabolism.

To further capture the patterns of expression changes in response to the SARS-CoV-2 infection, we performed a weighted co-expression network analysis on both transcriptomic and proteomic datasets using all genes and proteins detected, their functional assignments, and the top genes (Fig. 1f) and proteins (Fig. 1g) in key network elements. For transcriptomic and proteomic networks (adjusted p value <0.01, Spearman ρ > 0.83), we identified a set of five transcriptomics and four proteomics communities of strongly interconnected genes and proteins (Fig. 1h). These communities were also validated against random networks. Characterization of these communities again highlighted several pathways of interest including HIF-1, mTOR, and TNF signaling previously observed in our pairwise comparisons and time series analyses (Fig 1e). Ranking of all communities based on their centrality further identified those that display a higher number of central genes/proteins, i.e. communities that exhibit a larger number of associated genes/proteins and thus capture most coordinated expression changes and hence are predicted to robustly influence network behavior. The two most central communities (Fig. 1f and 1g) entail several genes associated with AKT1, SLC2A1 (HIF-1 signaling), RAF1 (MAPK signaling), SEC13 (mTOR signaling) and Caspase 8 (CASP8; TNF signaling). Functional enrichment analysis indicated that these two communities were associated (adjusted p< 0.05) with mTOR and MAPK signaling, Lysosomal and Proteasome-related processes, and cell cycle control. Importantly, we found that MAPK, AKT1, and mTOR showed cumulative expression changes through time (Fig. S1a and S2a); moreover, they were co-expressed (adjusted p<0.05) with several other genes/proteins, including those most central in each community (Fig. S1b,c and S2b,c), further highlighting the importance of these genes/proteins in coordinating the global response to infection. Finally, we found significant (adjusted p<0.05) functional overlap between three transcriptomic and three proteomic communities (communities 2, 3, 4; and communities A, B, D; Fig. 1h), thus pointing to common biological responses at the proteo-transcriptomic levels. These intersections included mTOR signaling (community 3 vs B), oxidative phosphorylation (communities 2,4 vs B) and thermogenesis (community 2,3,4 vs B), which are simultaneously found among similar communities in transcriptomic and proteomic networks. Our functional and network community analyses identify common host cell genes (AKT1, MAPK) and biological pathways (mTOR, MAPK and HIF-1 signaling) that are upregulated in SARS-CoV-2 infected HuH7 cells.

#### Dysregulated proteins and effector molecules in mTOR/HIF-1 signaling

The top four identified pathways, ErbB, PI3K-Akt, HIF-1, and mTOR signaling showed overlap of several proteins like AKT1, mTOR, MAPK, 4E-BP1, and S6K as represented in Sankey plot (Fig 2a). Since all the top identified pathways converge at mTOR signaling, we wanted to investigate whether SARS-CoV-2 infections indeed change expression of critical effector molecules of the mTOR/HIF-1 signaling pathway, namely 4E-BP1, S6K1 and HIF-1α. The mTOR pathway is involved in various biological functions and several viruses hijack this pathway to promote their own replication in different ways ^5^. To this end the phosphorylation status of the effector molecules of the mTOR signaling was assessed during SARS-CoV-2 infection. The infection dynamics measured by SARS-CoV-2 RNA in the cell culture supernatant and in the cell is shown in Fig 2b. Western blot results showed that the phosphorylation states of 4E-BP1 and S6K1 markedly changed during the course of the infection as compared to the mock infected control cells, showing a significant dip at 72 hpi (Fig 2c and 2d). Furthermore, HIF-1α protein levels were rapidly reduced following SARS-CoV-2 infections (Fig 2c and 2d).

**Figure 2.**
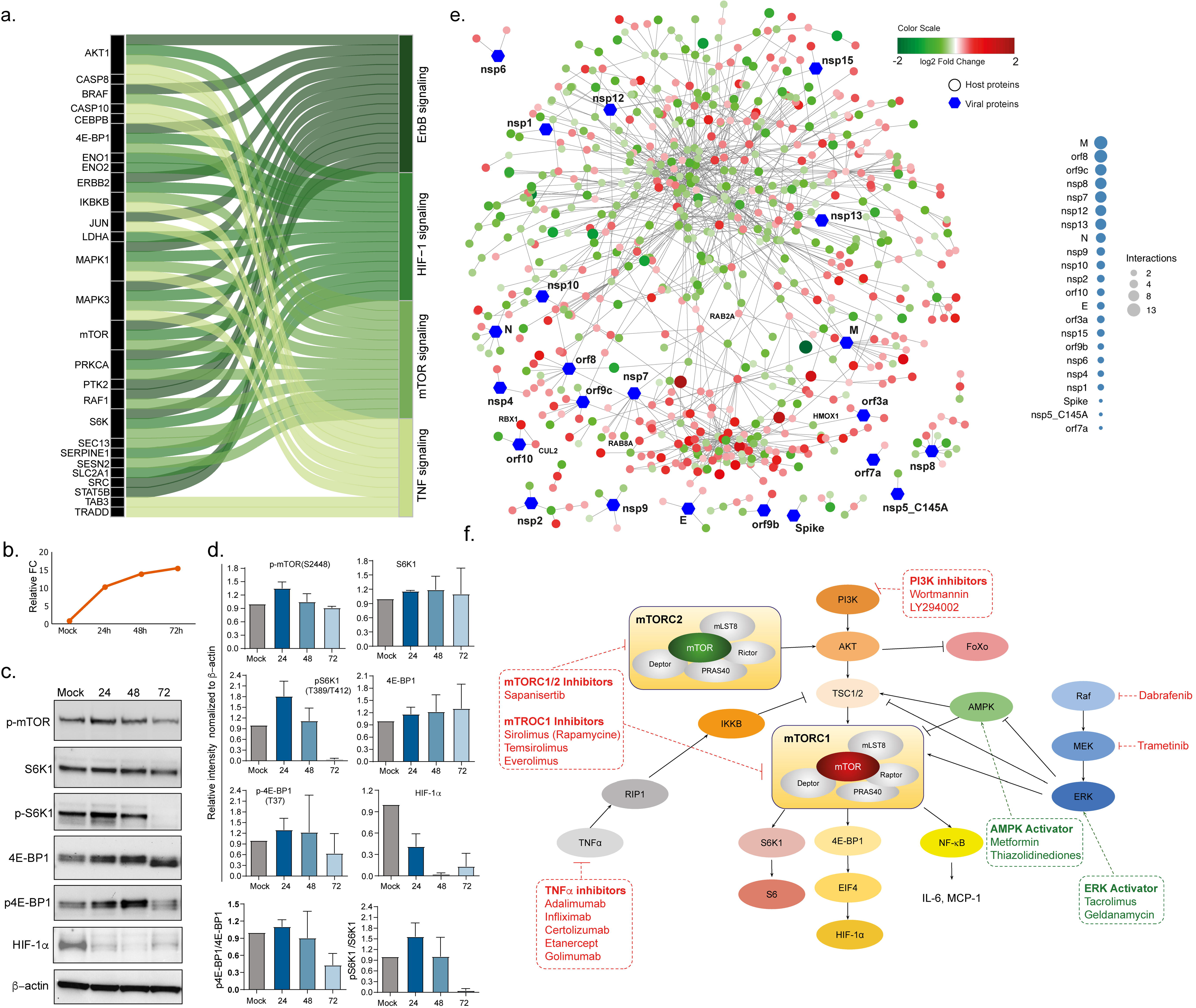
(**a**) Top four pathways, ErbB signaling, HIF-1 signaling, mTOR signaling and TNF signaling were selected and, together with proteins that are altered in the infection course, represented as Sankey Plot in order to illustrate the most important contribution to the flow of each pathway. **(b)** HuH7 cells were infected with SARS-CoV-2 at MOI of 1 and cells were harvested at 24hpi, 48hpi and 72hpi. The viral RNA quantification using qPCR targeting the E gene of SARS-CoV-2 targeting the supernatant. The relative fold change with respect to the uninfected controls is shown. **(c)** The representative western blots of indicated antibodies with **(d)** densitometric quantification are shown. **(e)** Network visualizing protein interactions among significantly changing proteins between samples at 24hpi and 48hpi, and SARS-CoV-2 viral proteins. Green color nodes represent decreased proteins at 48hpi and red colored proteins represent increased proteins at 48hpi. Size of the nodes are relative to their log2 fold change. Hexagonal shaped nodes denote SARS-CoV-2 viral proteins. The edges are derived from Human Reference Interactom (HuRI) and SARS-CoV-2 entry in Human Protein Atlas. **(f)** Approved drugs targeting AKT/mTOR/HIF-1 signaling pathway. Only key proteins of the pathway are shown. The inhibitors are shown in red and activators in green.

#### Drug repurpose and viral host protein interactions of the dysregulated proteins

To repurpose antiviral drugs targeting host-viral interactions is an attractive strategy to find drugs that might work against COVID-19. Therefore, we assessed protein-protein interactions including both host protein and SARS-CoV-2 protein associations obtained from the Human Protein Atlas (https://www.proteinatlas.org/humanproteome/sars-cov-2).^6^ Proteins that were significantly increased between 24h and 48h after SARS-CoV-2 infection (Fig 2e) were arbitrarily assigned to early responses. A total of 108 host-viral protein interactions were observed. The majority of the interactions was observed with the viral protein M (13 interactions), followed by orf8 (12 interactions), orf9c (11 interactions), nsp7, nsp8 (9 interactions) and nsp12 (8 interactions). Interestingly, while mapping these proteins with the pathway we observed that three HIF-1 pathway-associated proteins were markedly altered in the infected cells: Heme Oxygenase 1 (HMOX1), which interacts with orf3a, decreased over time, whereas Cullin 2 (CUL2) and Ring-Box 1 (RBX1), both of which interact with orf10, increased over time. In addition, we found that the two Ras-associated proteins RAB8A and RAB2A interact with nsp7. Furthermore, we found that Receptor-interacting serine/threonine-protein kinase 1 (*RIPK1*), involved in NF-κB, NLR, RIG-I-like receptor and TNF signaling,^7^ interacted with nsp12 and was significantly enriched over time (Fig 2e). Our data indicate a role of mTOR/HIF-1 in the cellular response to the SARS-CoV-2 infection, suggesting that drugs blocking this pathway could be possibly repurposed for COVID-19 patients (Fig 2f).

## Discussion

In this study using the integrated proteo-transcriptomics studies we identified four pathways, ErbB, HIF-1, mTOR and TNF signaling, among others that were markedly modulated during the course of the SARS-CoV-2 infection *in vitro*. Western blot validation of the downstream effector molecules of these pathways revealed a significant reduction in activated S6K1 and 4E-BP1 at 72 hours post infection. The data therefore points towards dysregulation of mTOR/HIF-1 signaling cascades, which could be potential target for COVID-19 therapeutic interventions.

The mTOR signaling pathways are known to regulate apoptosis, cell survival, and host transcription and translation and can be hijacked by several RNA viruses like influenza virus and coronaviruses.^8–11^ PI3K activation results in AKT phosphorylation and subsequent activation of mTOR. Through a cascade of events, mTORC1 and AKT activates 4E-BP1 and eIF4 complex followed by translation of effector protein HIF-1α that initiates host transcription and translation of specific genes. Another pathway that changed over time was the TNF signaling pathway. TNF signaling is also interlinked with HIF-1 signaling and can induce HIF-1α through AKT and MAPK activation.^12^ Of note, specific proteins dysregulated in the TNF signaling pathway were caspase 8, caspase 10,^13^ and CCAAT/enhancer-binding protein beta (CEBPB),^14^ which are linked to interferon (IFN) signaling and NF-κB signaling pathways. Previous studies on coronaviruses suggest a critical role of the IFN response, in particular IFN-β.^15,16^ This is also reflected in our findings that SARS-CoV-2 infections result in significantly dysregulated RIG-I, NLR and NF-κB pathways, which needs further evaluation. All these pathways have been linked to the IFN response.

It has been shown that absence of HIF-1α can promote replication of influenza A virus and severe inflammation mediated via promotion of autophagy.^17^ There could be several mechanisms that result in decreased phosphorylation of mTOR effectors such as the host response affecting the translation machinery in response to stress or viral proteins at the late stage of infection that promote the translation of viral mRNAs by shutting down host mRNA translation. Nonetheless, similar to other viruses that hijacking the AKT/mTOR pathway such as the highly pathogenic 1918 influenza virus^8^ and the Middle East respiratory syndrome coronavirus (MERS-CoV),^11^ deregulation of the mTOR pathway might enable SARS-CoV-2 to enhance its pathogenicity

A recent drug target network analysis based on potential human coronavirus and host interactions predicted that sirolimus (also known as rapamycin), which targets mTOR, could be repurposed.^18^ Sirolimus was shown to inhibit MERS-CoV infection by 60% in mice.^11^ Some studies have shown that everolimus, another mTOR inhibitor, and sirolimus are weakly active against influenza A virus.^19,20^ Everolimus delayed death but was not able to reduce mortality in lethal mouse infection model of influenza A (H1N1 and H5N1).^19^ Sirolimus was even shown to negatively affect the lung pathology probably due to its immunosuppressive effect.^20^ It has also been reported to block viral protein expression and virion release, improving the prognosis in patients with severe H1N1 pneumonia and acute respiratory failure.^21^ On the other hand, rapamycin treatment was shown to degrade antiviral barriers and could thus be potentially harmful in pathogenic viral infections.^22^ In COVID-19 patients the severity of the disease is associated with a cytokine storm with markedly increased expression of interleukin 6 (IL-6) in the serum of severe cases.^23^ Interestingly, IL-6 can activate mTOR in a STAT3 dependent or independent manner.^24^ Whether our proposed drugs can be indeed repurposed for COVID-19 therapies now needs to be carefully tested in *in vitro* SARS-CoV-2 infection models and in *in vivo* COVID-19 disease models.

There are some limitations of our study. First, we only used the HuH7 cell line but the SARS-CoV-2 can be also cultured in Vero E6, Vero CCL81, or HEK-293T cells. Whereas SARS-CoV-2 exerts rapid cytopathic effects in Vero E6 cells (within 24hpi), viral replication is slower in HuH7 and HEK293T cells, allowing to study host-cellular responses for 3 days after viral challenge ^25^. Moreover, an earlier study used HuH7 cells to identify the transcriptomics signature of early cellular responses to SARS-CoV and HCoV-229E infections ^26^. However, the observed effects could be cell type-specific and thus, we are currently assessing the effect of SARS-CoV-2 infection in other cells lines. Moreover, SARS-CoV-2 has a propensity to mutate and our experiments were performed with only one virus strain isolated from a Swedish patient. Of note, our virus isolate has close sequence similarity to the initial strains circulating in Wuhan, China.

In conclusions, we observed marked alterations of mTOR/HIF-1 signaling at the proteo-transcriptomic levels in response to SARS-CoV-2 infections, though the exact mechanistic role of these changes remains to be elucidated. Targeting mTOR/HIF-1 signaling could be an attractive candidate as a potential therapy, alone or preferably combined with antivirals, for the management of COVID-19 patients. Moreover, mTOR inhibition could be used to reduce the cytokine storm syndrome in severe cases of COVID-19.

## Methods

### Cells and viruses

The SARS-CoV-2 virus was isolated from a nasopharyngeal sample of a patient in Sweden and the isolated virus was confirmed as SARS-CoV-2 by sequencing (Genbank accession number MT093571). The hepatocyte derived cellular carcinoma cell line Huh7 was used. The cells were obtained from Marburg Virology Lab, Philipps-Universität Marburg, Marburg, Germany fully matching the STR reference profile of HuH-7. ^27^

### SARS-CoV-2 infection of Huh7 cells

Huh7 cells were plated in 6 well plates (2,5×10^5^ cells/well) in DMEM (Thermo Fisher Scientific, US) supplemented with 10% heat-inactivated FBS (Thermo Fisher, US). At 90-95% cell confluence the medium was removed, cells washed carefully with PBS and thereafter either cultured in medium only (uninfected control) or infected with SARS-CoV-2 at a multiplicity of infection (MOI) of 1 added in a total volume of 0.5 mL. After 1 hr of incubation (37°C, 5%CO_2_) the inoculum was removed, cells washed with PBS and 2 mL DMEM supplemented with 5% heat-inactivated FBS was added to each well. Samples were collected at three different time points, 24, 48 and 72 hrs post-infection (hpi). Samples were collected for proteomics and western blot, and RNAseq.

### Total RNA extraction and Quantification of viral RNA

The cells (uninfected, 24hpi, 48hpi and 72hpi) were collected by adding Trizol™ (Thermo Fisher Scientific, US) directly to the wells. RNA from SARS-CoV-2 infected and uninfected Huh7 cells and supernatent was extracted using the Direct-zol RNA Miniprep (Zymo Research, US) and quantitative real-time polymerase chain reaction (qRT-PCR) was conducted using TaqMan Fast Virus 1-Step Master Mix (Thermofisher Scientific, US) with primers and probe specific for the SARS-CoV-2 E gene following guidelines by the World Health Organization (https://www.who.int/docs/default-source/coronaviruse/wuhan-virus-assayv1991527e5122341d99287a1b17c111902.pdf) as described previously ^4^.

### Transcriptomics analysis (Illumina RNAseq)

The samples were sequenced using Illumina NextSeq550 in single-end mode with read length of 75 bases. The raw sequence data were first subjected to quality check using FastQC tool kit version 0.11.8. Illumina adapter sequences and low-quality bases were removed from the raw reads using the tool Trim Galore version 0.6.1. Phred score of 30 was used as cut-off to remove low quality bases. Quality of the data was again checked after pre-processing to assure high quality data for further analysis. The pre-processed reads were then aligned against human reference genome version 38 Ensembl release 96. Short read aligner STAR version 2.7.3a was used for the alignment. STAR was executed by setting the parameter soloStrand to Reverse to perform strand specific alignment and rest of the required parameters were set to default. The alignment result was written in sorted by co-ordinate bam format. After the alignment gene level read count data was generated for each sample using the module featureCounts from the software subread version 2.0.0. Read counting was performed by setting attribute type in the annotation to gene_id and strand specificity to reverse. Human reference gene annotation version 38 Ensembl release 96 in gtf format was used for the read counting. Normalization factors were calculated using the R package edgeR ^28^ from read counts matrix to scale the raw library sizes. Low expression genes with maximum counts per million (CPM) values under 1 per sample were removed from the sample. As recommended in RNAseq, data were transformed to CPM and variance weight was calculated using voom function. Square root of residual standard deviation against log2 CPMs was plotted to verify transformation quality.

### Protein extraction and in-solution digestion

The cells (uninfected, 24hpi, 48hpi and 72hpi) were lyzed in lysis buffer (5% glycerol, 10 mM Tris, 150 mM NaCl, 10% SDS and protease inhibitor), NuPAGE™ LDS sample buffer (ThermoFisher Scientific,US) was added and the samples was boiled at 99°C for 10 min. Aliquots of cell lysates (150 μL) were transferred to sample tubes and incubated at 37°C for 5 min at 550 rpm on a block heater and sonicated in water batch for 5 min. Each sample was reduced by adding 7 μL of 0.5M dithiothreitol (DTT) at 37°C for 30 min and alkylated with 14 μL of 0.5M iodoacetamide for 30 min at room temperature (RT) in the dark. Following addition of 2 μL of concentrated phosphoric acid and 1211 μL of binding buffer, protein capturing was performed according to the manufacturer’s protocol using S-Trap™ Micro spin columns (Protifi, Huntington, NY). After washing with 150 μL of binding buffer four times the samples were subjected to proteolytic digestion using 1.2 μg trypsin (sequencing grade, Promega) for 2h at 47°C. Then 40 μL of 50 mM TEAB was added following acidification with 40 μL of 0.2% formic acid (FA) and elution with 40 μL of 50% acetonitrile (AcN)/0.2% FA and the eluents were dried using a Vacufuge vacuum concentrator (Eppendorf, US). The resulted peptides were cleaned up in a HyperSep filter plate with bed volume of 40 ***μ***L (Thermo Fisher Scientific, Rockford, IL). Briefly, the plate was washed with 80% AcN/0.1% FA and equilibrated with 0.1% FA. Samples were filtered in the plate and washed with 0.1% FA. Peptides were eluted with 30% AcN/0.1% FA and 80% AcN/0.1% FA and dried in a vacuum concentrator prior to tandem mass tag (TMT) labeling.

### TMT-Pro labeling

Dry samples were dissolved in 30 μL of 100 mM triethylammonium-bicarbonate (TEAB), pH 8, and 100 μg of TMT-Pro reagents (Thermo Scientific,US) in 15 μL of dry acetonitrile (AcN) were added. Samples were scrambled and incubated at RT at 550 rpm for 2 h. The labeling reaction was stopped by adding 5 μL of 5% hydroxylamine and incubated at RT with 550 rpm for 15 min. Individual samples were combined to one analytical sample and dried in vacuum concentrator.

#### High pH reversed phase LC fractionation and RPLC-MS/MS analysis

The TMTPro-labeled tryptic peptides were dissolved in 90 μL of 20 mM ammonium hydroxide and were separated on an XBridge Peptide BEH C18 column (2.1◻mm inner diameter × 250◻mm, 3.5◻μm particle size, 300 Å pore size, Waters, Ireland) previously equilibrated with buffer A (20 mM NH_4_OH) using a linear gradient of 1–23.5% buffer B (20◻mM NH_4_OH in AcN, pH 10.0) in 42 min, 23.5%-54% B in 4 min and 54-63% B in 2◻min at a flow rate of 200◻μL/min. The chromatographic performance was monitored by sampling eluate with a UV detector (Ultimate 3000 UPLC, Thermo Scientific, US) monitoring at 214◻nm. Fractions were collected at 30 s intervals into a 96-well plate and combined into 12 samples concatenating eight fractions representing the peak peptide elution. Each combined fraction sample (800 μL) was dried in a vacuum concentrator and the peptides was resuspended in 2% AcN/0.1% FA prior to LC-MS/MS analysis.

Approximately, 2μg samples were injected in an Ultimate 3000 nano LC on-line coupled to an Orbitrap Fusion Lumos mass spectrometer (Thermo Scientific, San José, CA). The chromatographic separation of the peptides was achieved using a 50 cm long C18 Easy spray column (Thermo Scientific,US) at 55°C, with the following gradient: 4-26% of solvent B (2% AcN/0.1% FA) in 120 min, 26-95% in 5 min, and 95% of solvent B for 5 min at a flow rate of 300 nL/min. The MS acquisition method was comprised of one survey full mass spectrum ranging from *m/z* 350 to 1700, acquired with a resolution of R=120,000 (at *m/z* 200) targeting 4×10^5^ ions and 50 ms maximum injection time (max IT), followed by data-dependent HCD fragmentations of precursor ions with a charge state 2+ to 7+ for 2 s, using 60 s dynamic exclusion. The tandem mass spectra were acquired with a resolution of R=50,000, targeting 5×10^4^ ions and 86 ms max IT, setting isolation width to *m/z* 1.4 and normalized collision energy to 35% setting first mass at *m/z* 100.

### Peptide identification and preprocessing

The raw files were imported to Proteome Discoverer v2.4 (Thermo Scientific) and searched against the *Homo sapiens* SwissProt (2020_01 release with 20,595 entries) and the pre-leased SARS-CoV-2 UniProt (completed with 14 SARS-CoV2 sequences of COVID-19 UniProtKB release 2020_04_06) protein databases with Mascot v 2.5.1 search engine (MatrixScience Ltd., UK). Parameters were chosen to allow two missed cleavage sites for trypsin while the mass tolerance of precursor and HCD fragment ions was 10 ppm and 0.05 Da, respectively. Carbamidomethylation of cysteine (+57.021 Da) was specified as a fixed modification, whereas TMTPro at peptide N-terminus and lysine, oxidation of methionine (+15.995 Da), deamidation of asparagine and glutamine were defined as variable modifications. For quantification both unique and razor peptides were requested. Protein raw data abundance was first filtered for empty rows with *in house* script and quantile-normalize using R package NormalyzerDE ^29^. Principal component analysis (PCA) was applied to explore sample-to-sample relationships. One proteomics samples from the uninfected control was excluded as it turned out to be outlier.

### Statistical analysis

Proteomics and transformed transcriptomics data were tested for normality using histograms with normal distribution superimposed. Differential expression through linear model was performed using R package LIMMA ^30^. LIMMA supports multifactor designed experiments in microarray, transcriptomics and proteomics. Its features are designed to support small number of arrays. The three infected replicates at 24hpi, 48hpi and 72hpi hours respectively were selected in order to perform an equi-spaced univariate time series analysis. In limma design matrix, separated coefficients were associated with time and replicates in order to extract the difference as a contrast. Moderated paired-t-test using limma with adjustment for replicates was applied. For pairwise comparisons, single factorial design was implemented to fit model with a coefficient for each of our four factors: uninfected, 24hpi, 48hpi and 72hpi. Comparisons were extracted as contrasts. In both analysis, significant differential genes and proteins were selected based on p values after Benjamini-Hochberg (BH) adjustment. Genes with alpha value inferior to 0.05 were considered significant.

#### Bioinformatics Analysis

The transcriptomics and proteomics analysis were performed using all the protein coding genes and proteins and a gene set of viral processes, response and diseases respectively. The viral response gene set is a catalogue of genes which are known to have involve in viral processes, response and diseases. The catalogue was enriched by mining biological process category of gene-ontology terms, Reactome pathways and gene sets associated with various viral diseases. Gene Ontology terms were selected by keeping, “response to virus (GO:0009615)” as parent term. All child terms of GO:0009615 were selected based on ontology term relationship “is a” and “regulates”. The pathway “Antiviral mechanism by IFN-stimulated genes” and two other events it participates were selected from Reactome database. Gene sets related to 42 virus associated diseases and six virus related diseases were selected from “Rare_Diseases_AutoRIF_Gene_Lists” library provided by gene set enrichment tool Enrichr ^31^. The viral response gene set contains total of 1517 protein coding genes. After filtering antiviral genes, up and downregulated proteins and transcripts were submitted separately to gene set enrichment analysis (GSEA) using gseapy v0.9.17.

R package gplots v3.03 was used to generate heatmaps to display terms associated adjusted p values contrasts over conditions.

### Network and community analyses

Association analyses were performed by computing pairwise Spearman rank correlations for all features after removing null variant or genes with very low expression (RPKM < 1). Correlations were considered statistically significant at FDR < 0.01. Positive correlations were selected and used to build a weighted graph where Spearman ρ was used as weights. All network analyses were performed in igraph ^32^. For all networks, diameter, average path lengths, clustering coefficients, and degree distributions were compared with those attained for similarly-sized random networks (Erdős-Rényi models, ^33^). Degree centrality was computed for all networks and normalized for network size. Communities were identified by modularity maximization through the Leiden algorithm ^34^. Community centrality was computed by averaging node centrality and used to identify the most central communities in each network by degree comparison. Gene set enrichment analysis was performed on each community (n > 30) through Enrichr for KEGG Human 2019 where backgrounds were selected based on the node number of each network. Community similarity was computed through hypergeometric testing of overlap between statistically significant KEGG terms for each transcriptomic vs proteomic pair of communities. Throughout, all statistical tests were considered at an FDR < 0.05 unless otherwise stated. All analyses were performed in Python 3.7.

Protein-protein interactions among human proteins were derived from Human Reference Interactome (HuRI). Interactions between human proteins and SARS-Cov2 viral proteins were obtained from Human Protein Atlas (HPA). Protein interaction network is created using Cytoscape version 3.6.1 ^35^. Edge weighted spring embedded layout was used for the network. R package gplots 3.03 was used to generate heatmaps to display terms associated p values contrasts over conditions. Sankey Plot illustrates most important contribution genes to flow pathways. It was plotted using R package ggalluvial version 0.11.1 ^36^. Scatter plots produced using ggplot2 represent the bivariate relationship between proteins and time.

### Western Blot

Evaluation of protein expression was performed by running 20 μg of total protein lysate on NuPage Bis Tris 4-12%, gels or NuPage Tris-Acetate 3-8% gels (Invitrogen, Carlsbad, CA, USA). Proteins were transferred using iBlot dry transfer system (Invitrogen, Carlsbad, CA, USA) and blocked for 1h using 5% milk or bovine serum albumin (BSA) in 0.1% PBSt (0.1% Tween-20). Subsequent antibody detection was performed at 4°C over-night or 2h at room temperature for β-Actin. Membranes were washed using 0.1% PBSt and secondary antibody incubated 1h at room-temperature using Dako Polyconal Goat Anti-Rabbit or Anti-Mouse Immunoglobulins/HRP (Aglient Technologies, Santa Clara, CA, USA) washed using 0.1% PBSt and visualized using ECL or ECL Select (GE Healthcare, Chicago, IL, USA) on ChemiDoc XRS+ System (Bio-Rad Laboratories, Hercules, CA, USA). The western blot analysis was performed on duplicates (p-mTOR, S6K and p-S6K) or triplicate (4E-BP1, p-4E-BP1and HIF-1α) of the samples in two different timepoints. Viral RNA was quantifed from cells as well as supernatent in all the time points as a confirmation of the infection. The uncropped western blots are given as resource data file 1.

## Data and Code Availability

The raw RNAseq data can be obtainred from the SRA using the project id. PRJNA627100. Proteomics data can be obtained from https://zenodo.org/record/3754719#.XqgnSy2B3OQ. All the codes are available at github: https://github.com/neogilab/COVID19

## Acknowledgements

The study is funded by the Swedish Research Council Grant (2017-01330, UN). A. M. is supported by the Swedish research Council 2018-05766 and 2017-03126. J.M.P. is supported by the Canada 150 Research Chair program and CIHR rapid Response COVID-19 grant. We would like to thank the core facility Bioinformatics and Expression Analysis (BEA), supported by the board of research at the Karolinska Institute and the research committee at the Karolinska hospital specifically Fredrik Fagerström-Billai and Carolina Bonilla Karlsson for rapidly performing the RNAseq run. We also like to thank Proteomics Biomedicum; Karolinska Institute, Solna, and National Bioinformatics Infrastructure Support, Sweden for supporting the project. The computations were performed on resources provided by SNIC through Uppsala Multidisciplinary Center for Advanced Computational Science (UPPMAX) under Project SNIC2017-550 Authors also like to thank Beatriz Sá Vinhas and Elisa Saccon for continuous discussion and performing the literature review for the article.

## Authors Contributions

S.A. performed all the infection experiments, S.G., S.S.A., M.S. and S.K. performed the other lab experiments and wrote the method and result part of the manuscript, A.T.A., F.M., K.S. and R.B. performed the bioinformatics analysis and wrote the method and result part of the manuscript. Á.V. and M.S. performed the mass-spectrometry and wrote the method part of the manuscript, J.M.P. and A.M contributed with resource and manuscript writing and revision. S.G. and U.N. wrote the first draft of the manuscript reviewed by J.M.P., A.M. U.N. conceived and designed the study and contributed with resource. All authors approved the final version of the manuscript.

## Supplementary Materials

**Figure S1** – Gene expression (A) and co-expression (B – C) among key genes and top correlated and central genes in each community identified based on a transcriptomic network (communities 1-5). For each community we identified selected the top 10 genes (gray labels), ranked by their median centrality (median ranked degree, betweenness, closeness and eccentricity centralities), among the top 10% correlated gene in each community. Key proteins, previously associated with HIF-1a, mTOR, MAPK signaling and other top pathways, are highlighted in black (Fig. 1e). Spearman rank correlations were computed for all genes (B) and excluded if not statistically significant (C, FDR < 0.01). S6K-A3, S6K-B2, and 4E-BP1 respectively indicate genes RPS6KA3, RPS6KB2, and EIF4EBP1.

**Figure S2** – Protein abundance (A) and correlations (B – C) among key proteins and top correlated and central proteins in each community identified based on a proteomic network (communities A-D). For each community we identified selected the top 10 proteins (gray labels), ranked by their median centrality (median ranked degree, betweenness, closeness and eccentricity centralities), among the top 10% correlated proteins in each community. Key proteins, previously associated with HIF-1a, mTOR, MAPK signaling and other top pathways, are highlighted in black (Fig. 1e). Spearman rank correlations were computed for all proteins (B) and excluded if not statistically significant (C, FDR < 0.01). S6K-A3, S6K-B2, and 4E-BP1 respectively indicate genes RPS6KA3, RPS6KB2, and EIF4EBP1. Note that 4E-BP1 is among the top 10% most correlated genes in community B.

**Supplementary Source Data 1.** Original western blot.

