## Supplemental Figure S1 and S2 and Data Source File for "Dysregulation in mTOR/HIF-1 signaling identified by proteo-transcriptomics of SARS-CoV-2 infected cells"

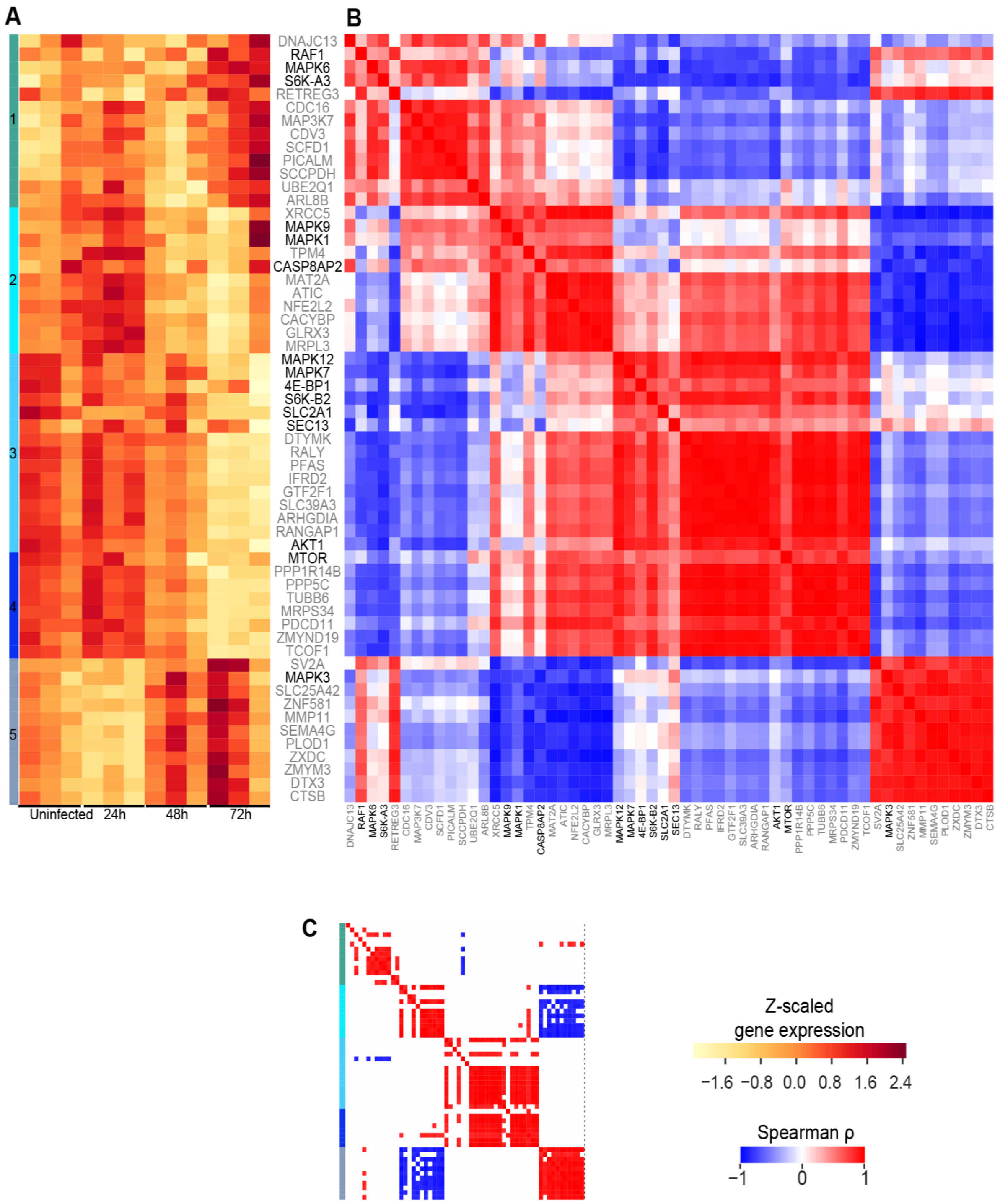


**Figure S1** – Gene expression (A) and co-expression (B – C) among key genes and top correlated and central genes in each community identified based on a transcriptomic network (communities 1-5). For each community we identified selected the top 10 genes (gray labels), ranked by their median centrality (median ranked degree, betweenness, closeness and eccentricity centralities), among the top 10% correlated gene in each community. Key proteins, previously associated with HIF-1a, mTOR, MAPK signaling and other top pathways, are highlighted in black (Fig. 1e). Spearman rank correlations were computed for all genes (B) and excluded if not statistically significant (C, FDR < 0.01). S6K-A3, S6K-B2, and 4E-BP1 respectively indicate genes RPS6KA3, RPS6KB2, and EIF4EBP1.

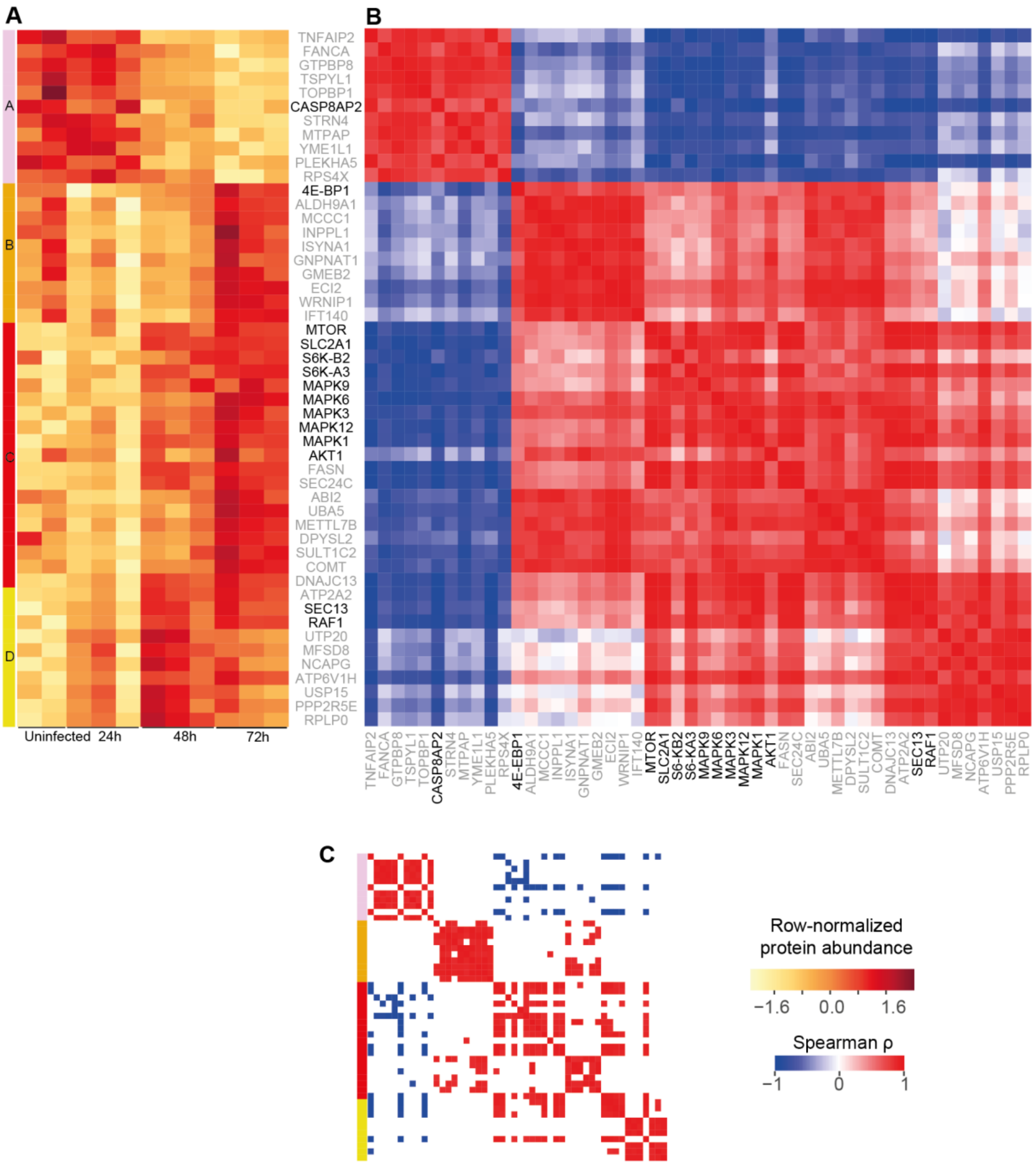


**Figure S3** – Protein abundance (A) and correlations (B – C) among key proteins and top correlated and central proteins in each community identified based on a proteomic network (communities A-D). For each community we identified selected the top 10 proteins (gray labels), ranked by their median centrality (median ranked degree, betweenness, closeness and eccentricity centralities), among the top 10% correlated proteins in each community. Key proteins, previously associated with HIF-1a, mTOR, MAPK signaling and other top pathways, are highlighted in black (Fig. 1e). Spearman rank correlations were computed for all proteins (B) and excluded if not statistically significant (C, FDR < 0.01). S6K-A3, S6K-B2, and 4E-BP1 respectively indicate genes RPS6KA3, RPS6KB2, and EIF4EBP1. Note that 4E-BP1 is among the top 10% most correlated genes in community B.

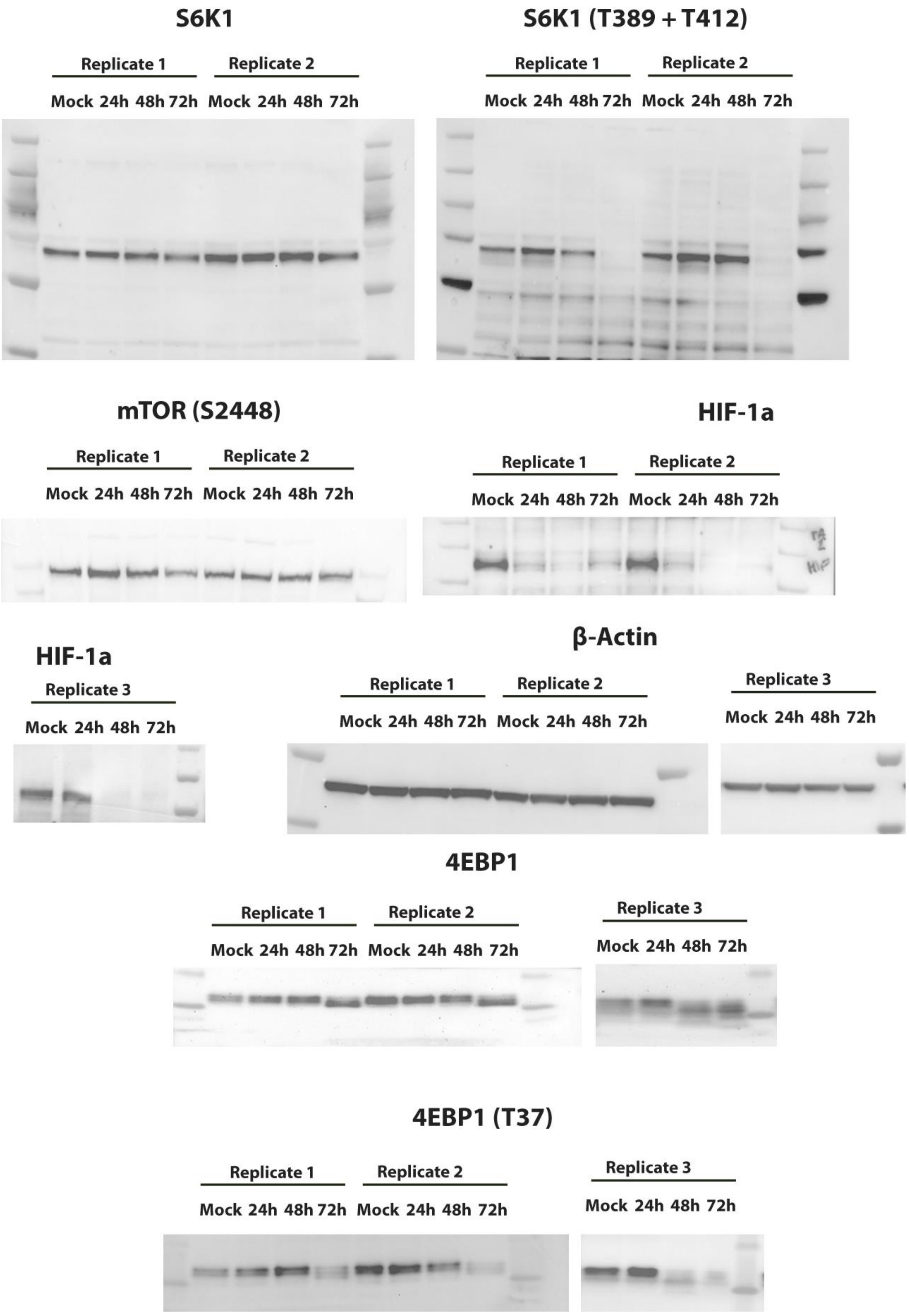


**Supplementary Source Data 1.** Original western blot.
